# iCn3D: From Web-based 3D Viewer to Structural Analysis Tool in Batch Mode

**DOI:** 10.1101/2021.09.10.459868

**Authors:** Jiyao Wang, Philippe Youkharibache, Aron Marchler-Bauer, Christopher Lanczycki, Dachuan Zhang, Shennan Lu, Thomas Madej, Gabriele H. Marchler, Tiejun Cheng, Li Chuin Chong, Sarah Zhao, Kevin Yang, Jack Lin, Zhiyu Cheng, Rachel Dunn, Sridhar Acharya Malkaram, Chin-Hsien Tai, David Enoma, Ben Busby, Nicholas L. Johnson, Francesco Tabaro, Guangfeng Song, Yuchen Ge

## Abstract

iCn3D was initially developed as a web-based 3D molecular viewer. It then evolved from visualization into a full-featured interactive structural analysis software. It became a collaborative research instrument through the sharing of permanent, shortened URLs that encapsulate not only annotated visual molecular scenes, but also all underlying data and analysis scripts in a FAIR manner. More recently, with the growth of structural databases, the need to analyze large structural datasets systematically led us to use Python scripts and convert the code to be used in Node.js scripts. We showed a few examples of Python scripts at https://github.com/ncbi/icn3d/tree/master/icn3dpython to export secondary structures or PNG images from iCn3D. Users just need to replace the URL in the Python scripts to export other annotations from iCn3D. Furthermore, any interactive iCn3D feature can be converted into a Node.js script to be run in batch mode, enabling an interactive analysis performed on one or a handful of protein complexes to be scaled up to analysis features of large ensembles of structures. Currently available Node.js analysis scripts examples are available at https://github.com/ncbi/icn3d/tree/master/icn3dnode. This development will enable ensemble analyses on growing structural databases such as AlphaFold or RoseTTAFold on one hand and Electron Microscopy on the other. In this paper, we also review new features such as DelPhi electrostatic potential, 3D view of mutations, alignment of multiple chains, assembly of multiple structures by realignment, dynamic symmetry calculation, 2D cartoons at different levels, interactive contact maps, and use of iCn3D in Jupyter Notebook as described at https://pypi.org/project/icn3dpy.

## 1 Introduction

With the release of more and more AlphaFold [1] or RoseTTAFold [2] predicted structures and experimentally determined structures, it is important to be able to do structural analysis in batch mode. Our recent development on iCn3D helps fill this need.

iCn3D is not only a web-based 3D structure viewer, but also an analysis tool either interactively or in batch mode. All interactive structural analyses in iCn3D could be converted to analyses in batch mode using either Python scripts or Node.js scripts, which call functions in iCn3D. This feature has been available since we modernized the code of iCn3D to use JavaScript classes. As shown in Figure 1, all recent additions are colored in orange. “Users” could use iCn3D for 3D views, annotations, or structural analyses through three different “Front End” interfaces: web pages, Jupyter Notebook, or Python/Node.js scripts in batch mode. The backends are “RESTful APIs” to retrieve 3D coordinates, annotations, or analysis data.

**Figure 1.**
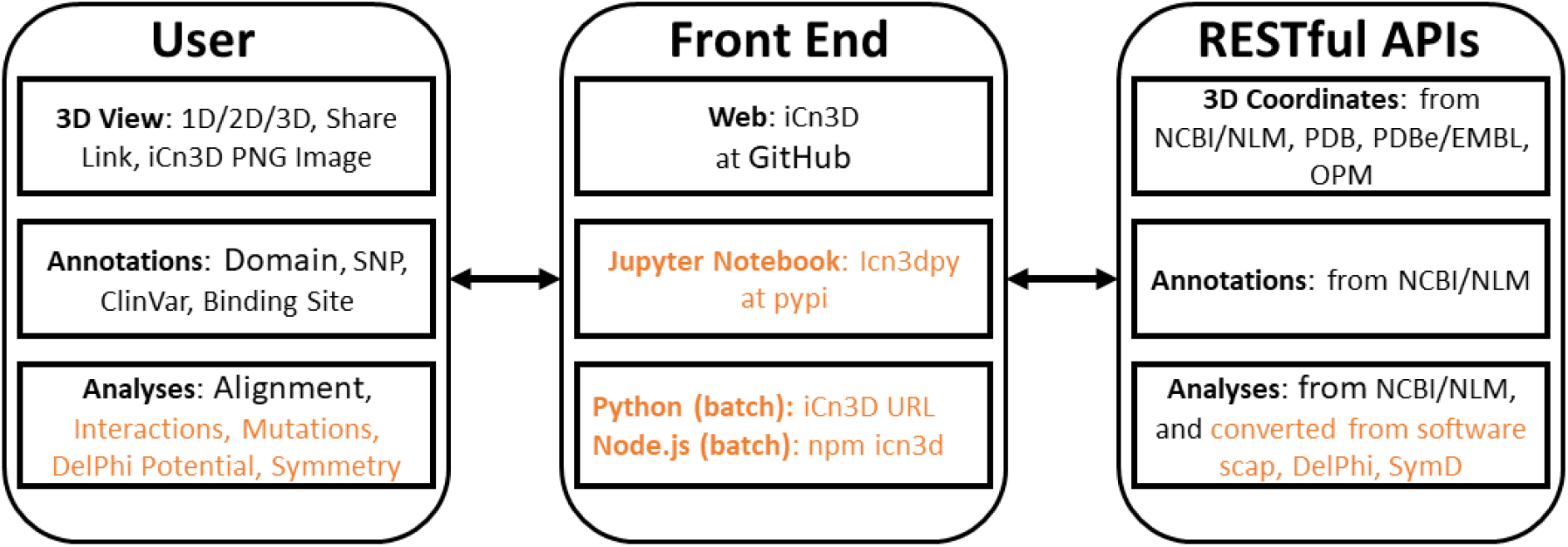
Front end and backend of iCn3D. The new additions are colored in orange.

iCn3D was first released in 2016 as a web-based 3D structure viewer with synchronized 1D, 2D and 3D views and many annotations. Any custom view from iCn3D can be shared in a permanent, shortened URL[3]. Since then, we introduced interaction analyses [4] and converted three pieces of software into RESTful APIs. First, DelPhi [5] was licensed from Columbia University and was used to calculate electrostatic potentials for nucleotides, proteins, and ligands in iCn3D. Second, scap [6] was kindly provided by the Honig group at Columbia University and was used to predict the side chain change due to mutation. Third, SymD [7] was kindly provided by Chin-Hsien Tai at the National Cancer Institute and was used to calculate protein symmetries dynamically [8].

We also added other features including the alignment of multiple chains from different structures, realignment, dynamic symmetry calculation for any subsets, 2D cartoons at different levels, interactive contact maps, and the use of iCn3D in Jupyter Notebook.

## Material and Methods

### 2.1 RESTful API based on DelPhi

The “DelPhi” program calculates electrostatic potentials. The C++ code of DelPhi [5] was licensed from Columbia University and converted to a RESTful API “delphi.cgi” at National Center for Biotechnology Information (NCBI). To qualitatively illustrate the electrostatic potential and speed up the calculation, the linear Poisson-Boltzmann equation was solved.

DelPhiPKa [9] is used to add hydrogens and partial charges to proteins and nucleotides. Open Babel [10] is used to add hydrogens to ligands. Antechamber [11] is used to add partial charges to ligands with the Gasteiger charge calculation method. The default grid size (n=65) and default salt concentration (0.15 M) can be changed. The pH is always set at 7.0. The parameters of all RESTful APIs are described at https://www.ncbi.nlm.nih.gov/Structure/icn3d/icn3d.html#restfulapi.

### 2.2 RESTful API based on scap

The “scap” program predicts side chain changes due to mutations. The C++ code of Jackal/scap [6] was provided by the Honig group and converted to a RESTful API “scap.cgi” at NCBI. The scap program first iteratively samples all sidechain rotamers until convergence. The final lowest-energy conformation among all the conformations starting from the initial 3 conformations will then be minimized by refining the side-chain conformation with 2-degree rotation on each bond in the sidechain to search for lower energy conformations around the rotamer. The large rotamer library was used.

### 2.3 RESTful API based on SymD

The “SymD” program determines symmetries in structures. The C++ code of SymD [7] was provided by the Chin-Hsien Tai and Byungkook Lee group and converted to a RESTful API “symd.cgi” at NCBI. The maximum allowed number of atoms is 30,000. A symmetry is considered found if the Z score is greater than 8. Only C_n_ and helical symmetries are considered.

### 2.4 Modernize iCn3D Code and Update Three.js

Before iCn3D version 3, two JavaScript objects “iCn3DUI” and “iCn3D” were used to hold all functions. In iCn3D version 3, we switched from ECMAScript 2009 (ES5) to ECMAScript 2015 (ES6) and used JavaScript classes. Each class is in a separate file and can access the classes “iCn3DUI” and “iCn3D”, which can access most functions. The class structure is described at https://www.ncbi.nlm.nih.gov/Structure/icn3d/icn3d.html#classstructure.

We also updated Three.js, which is used to generate and view 3D objects, from version 103 to version 128, which uses JavaScript classes as well. Since the changes on both iCn3D and Three.js are significant and JavaScript classes do not work in the Internet Explorer, we kept the previous library names untouched, and used new library names for Three.js and iCn3D. The updated embedding method is described at https://www.ncbi.nlm.nih.gov/Structure/icn3d/icn3d.html#HowToUse.

### 2.5 Build iCn3D Source Code

All JavaScript classes are bundled with rollup (https://rollupjs.org/guide/en/). Then gulp (https://gulpjs.com) was used to set up all files for release in iCn3D GitHub page (https://github.com/ncbi/icn3d) and icn3d npm package (https://www.npmjs.com/package/icn3d).

## 3 Results

### 3.1 Interaction Analysis

iCn3D shows eight kinds of interactions between molecular structures: contact, hydrogen bond, salt bridge or ionic interaction, halogen bond, π-cation interaction, and π-stacking. Users can generate the interactions shown in Figure 2 in three steps: click the menu “Analysis > Interactions”, select the two sets chain 6M0J_A and 6M0J_E in the popup window, and click “2D Interaction Network”. Figure 2A shows the interacting residues between ACE2 (in pink) and SARS-CoV-2 spike protein (in blue) using the PDB structure 6M0J. The green, cyan, and gray dotted lines show the hydrogen bonds, salt bridges, and contacts, respectively. Figure 2B shows the 2D interaction network, where the nodes represent residues, and the lines represent interactions. The interaction includes multiple hydrogen bonds, two salt bridges and many contacts. The custom view can be reproduced with a permanent sharable link https://structure.ncbi.nlm.nih.gov/icn3d/share.html?oLwZzGL59izJeEBVA as described in Figure 2.

**Figure 2.**
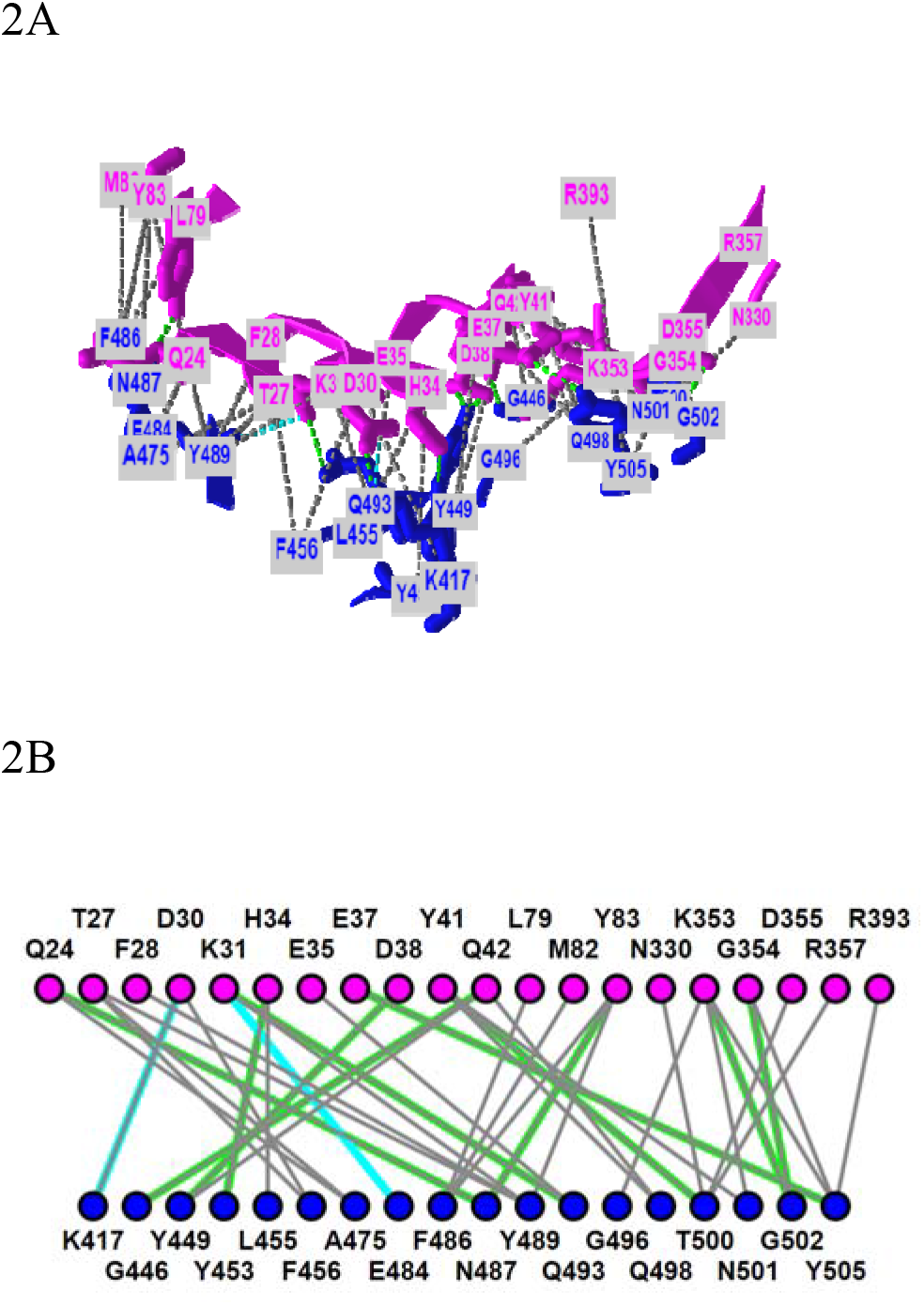
3D and 2D views of the interaction between ACE2 (pink) and SARS-CoV-2 spike protein (blue) (PDB 6M0J). (A) 3D view of the interaction. Hydrogen bonds, salt bridges, and contacts are depicted as dotted lines and are colored in green, cyan, and gray, respectively. (B) 2D interaction network of the interaction. Each node and line can be clicked to show the corresponding residue and interaction in the 3D view. Sharable Link: https://structure.ncbi.nlm.nih.gov/icn3d/share.html?oLwZzGL59izJeEBVA

This interactive analysis can be converted to a Node.js script https://github.com/ncbi/icn3d/blob/master/icn3dnode/epitope.js by using the npm package icn3d (https://www.npmjs.com/package/icn3d). Users can simply run the command “<u>node epitope.js</u> *<u>6M0J E A</u>*” in a command line to get the interacting residues between the chain *E* (spike protein) and chain *A* (ACE2) of the PDB structure *6M0J*.

### 3.2 Mutation Analysis

iCn3D can show residue mutations in 3D views. The side chain prediction is done with a RESTful API based on scap [6]. Users can show the change of interactions due to mutations by clicking the menu “Analysis > Mutation”, inputting the mutation (such as “*6M0J*_*E*_*501*_*Y*” to indicate the mutation is at position *501* of the *E* chain of PDB *6M0J* and the mutant is *Y*/Tyr residue), and clicking the button “Interactions”. Users can press the key “a” to toggle between the wild type and the mutant. Figure 3A shows the mutant’s 3D structure in the stick style. The spike protein is in blue and ACE2 is in pink. Figure 3B shows that the mutant Y501 (bottom panel) introduces one π-cation interaction (red line) and one π-stacking (blue line). The view can be reproduced with the sharable link as described in Figure 3. The interactive analysis can be converted to a Node.js script https://github.com/ncbi/icn3d/blob/master/icn3dnode/interaction2.js. Users can run the command “<u>node interaction2.js 6M0J E 501 Y</u>” to get the change of interactions.

**Figure 3.**
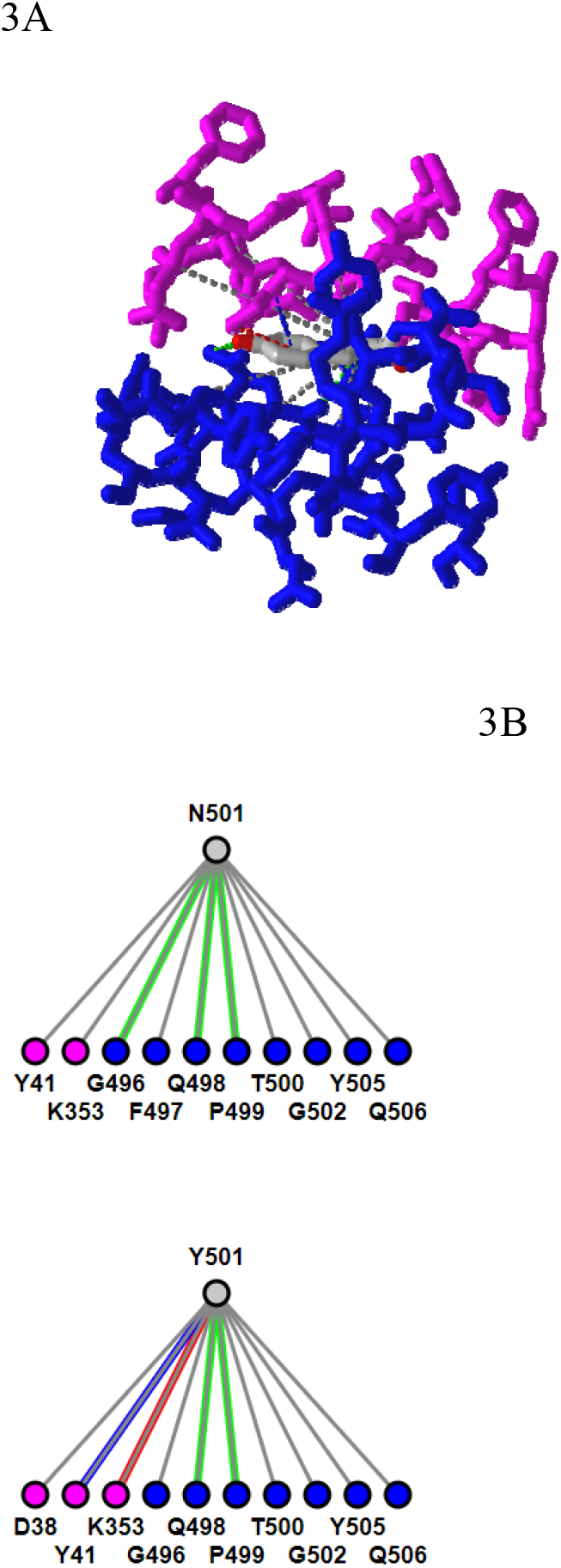
3D and 2D views of mutation N501Y of SARS-Cov-2 spike protein (PDB 6M0J). (A) 3D view of the mutant with Asn at position 501 mutated to Tyr. The spike protein is colored in blue and ACE2 is colored in pink. Hydrogen bonds, contacts, π-cation interaction, and π-stacking are depicted as dotted lines and are colored in green, gray, red, blue, respectively. (B) Change of interactions due to mutation. The top panel shows the wild type, and the bottom panel shows the mutant. Sharable Link: https://structure.ncbi.nlm.nih.gov/icn3d/share.html?j5Y5CkMQPU2R6smf9

### 3.3 DelPhi Electrostatic Potential

iCn3D can also show electrostatic potential for 3D structures. The electrostatic potential is calculated with a RESTful service based on the software DelPhi [5], which is licensed from Columbia University to be used in iCn3D. Figure 4A shows the binding of the PH domain of phospholipase C Delta to IP_3_ (or a membrane containing PIP_2_). The binding region contains multiple positively charged residues (in blue). Figure 4B shows the electrostatic potential on the surface of the PH domain excluding IP_3_. Blue and red indicate greater than +50 mV and less than −50 mV, respectively. The binding region clearly has high positive electrostatic potential. The custom view of Figure 4B can be reproduced with a sharable link: https://structure.ncbi.nlm.nih.gov/icn3d/share.html?5hvduy8LQHtJ1NdL6. The DelPhi electrostatic potential on the surface of chain A can be exported from a Node.js script https://github.com/ncbi/icn3d/blob/master/icn3dnode/delphipot.js with the command “node delphipot.js 1MAI A”. Users could also choose to show equipotential maps (Figure 4C) instead of surface potentials for the pH domain excluding IP_3_. The blue and red mesh show the +50 mV and −50 mV equipotential profiles.

**Figure 4.**
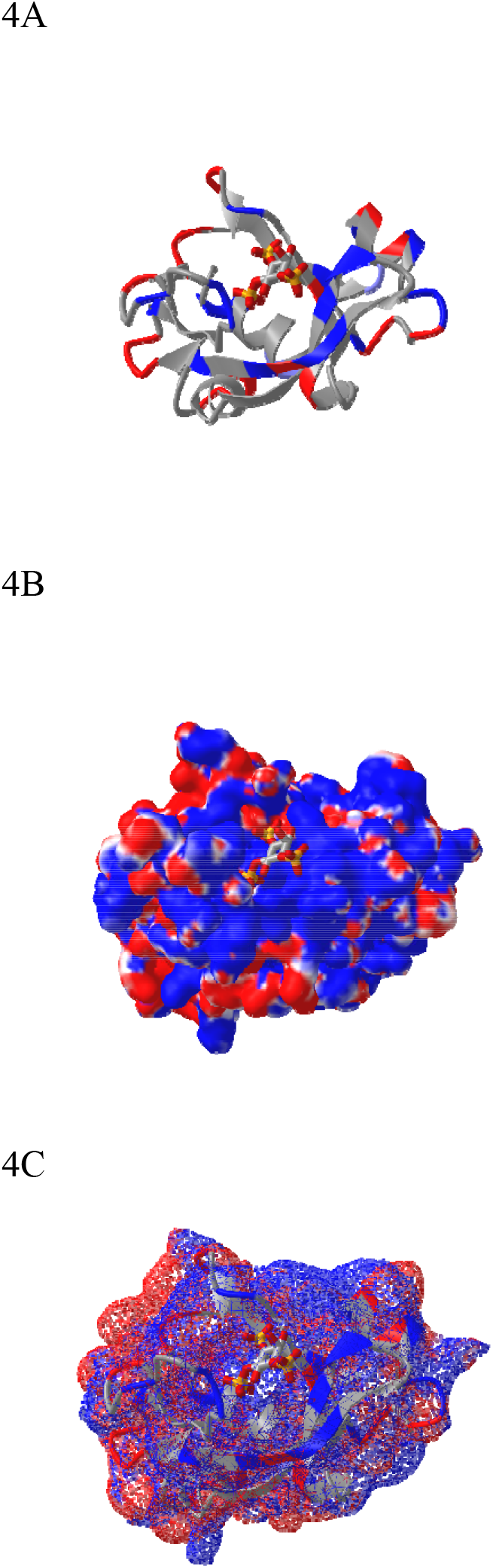
DelPhi electrostatic potential of membrane-binding PH domain (PDB 1MAI). (A) The PH domain is shown in ribbon style with positively and negatively charged residues colored in blue and red respectively. IP_3_ is shown in stick style with three phosphate groups. (B) The electrostatic potential of the PH domain is shown on the surface. Blue indicates greater than +50mV and red indicates less than −50 mV. Sharable link: https://structure.ncbi.nlm.nih.gov/icn3d/share.html?5hvduy8LQHtJ1NdL6. (C) The equipotential map of the PH domain. The blue and red meshes indicate +50 and −50 mV potential profile, respectively.

### 3.4 Dynamic Symmetry Calculation

iCn3D can not only show the pre-calculated symmetry from PDB, but also calculate symmetry for any selected residues. The calculation is done with a RESTful API based on the software SymD [7]. Figure 5B shows the structure of PDB 3HUJ. Even though the whole structure has no symmetry, its subset of residues 1-178 in chain A (colored in blue and red) has C2 symmetry as shown in Figure 5A. The red and blue colors indicate same residues and different residues in the symmetry alignment, respectively. The feature is available in the menu “Analysis > Symmetry > from SymD (Dynamic)”. The custom view can be reproduced in a sharable link as described in Figure 5.

**Figure 5.**
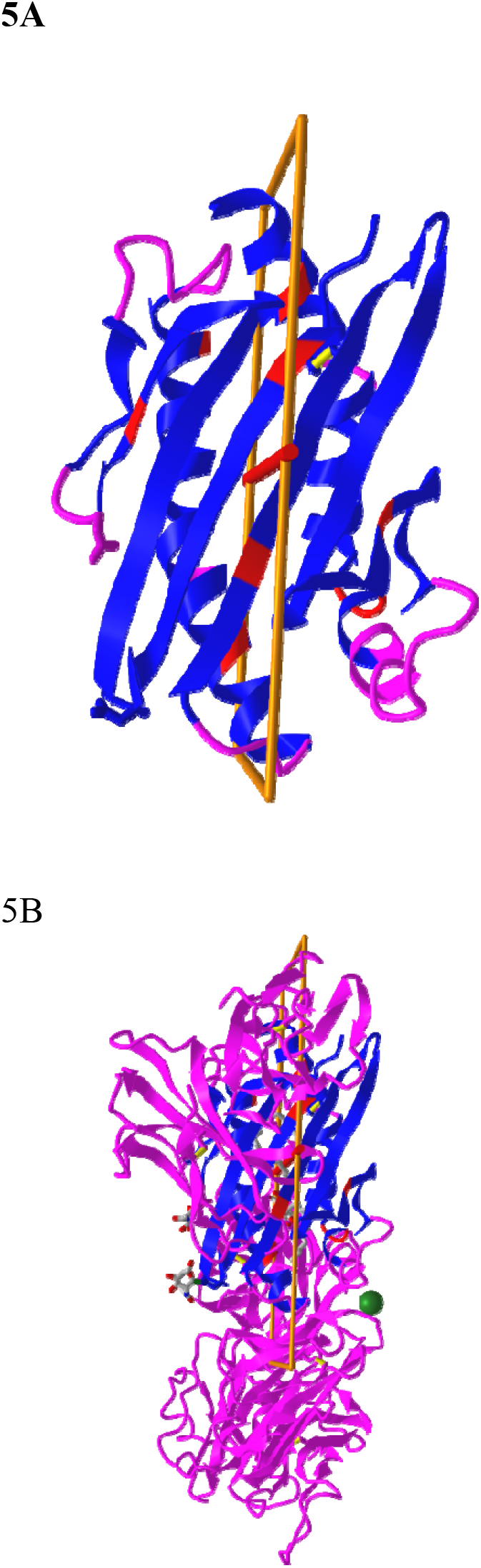
Dynamic symmetry calculation. (A) The subset of residues 1-178 in the A chain of PDB 3HUJ was used for symmetry calculation. The symmetric parts are colored in red (same residue) or blue (different residues). The rest are colored in pink. (B) The full structure of PDB 3HUJ with the symmetry in Chain A shown. Sharable link: https://structure.ncbi.nlm.nih.gov/icn3d/share.html?gNe7MemEb5vqYSba6.

### 3.5 Multiple Chain Alignment, and realignment

iCn3D can align multiple chains from different structures as shown in Figure 6 with two structures. The master structure is the first PDB structure 1HHO. All other structures are aligned to it. Figure 6A shows the alignment with full chains using the VAST algorithm [12] based purely on geometric criteria. The aligned residues are colored in red. Figure 6B shows the alignment to a subset of residue numbers 10-50 in the master structure with the following steps. First, a sequence alignment to the set of residues is done. Then the coordinates of these residues are used for a structure superposition. Figure 6C shows the alignment of predefined residue numbers 50-100 in each structure. The coordinates of the pre-defined residues are used for a structure superposition. The sharable links are shown in Figure 6.

**Figure 6.**
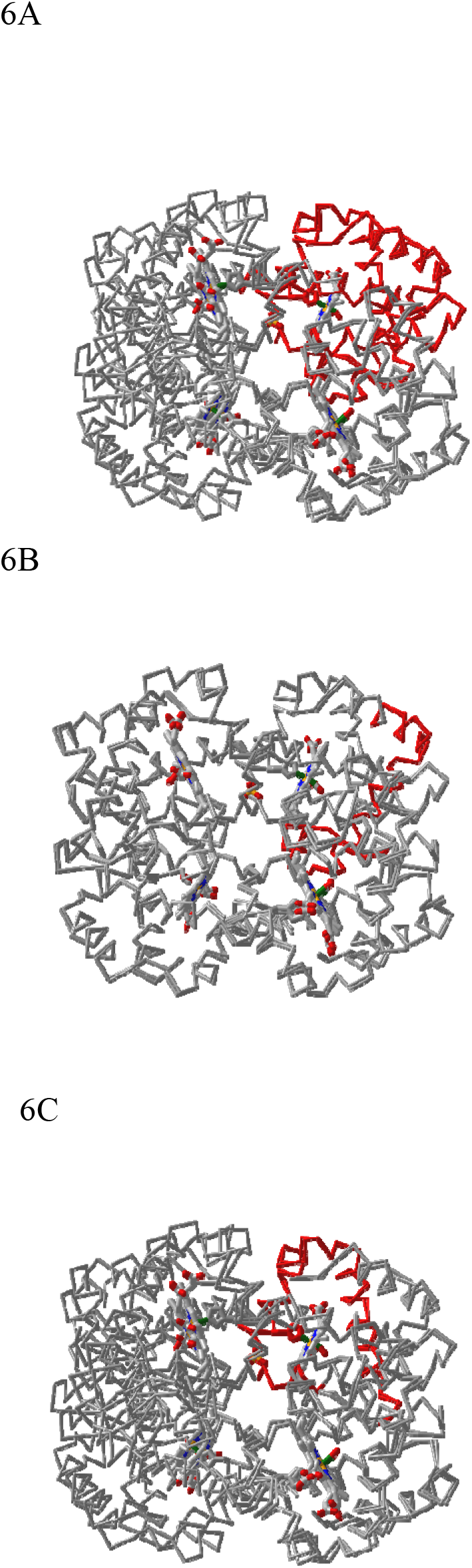
Multiple chain alignment of chain A of PDB 4N7N with chain A of PDB 1HHO. (A) Align with the full chains. The aligned part is colored in red. The sharable link is https://www.ncbi.nlm.nih.gov/Structure/icn3d/full.html?chainalign=1HHO_A,4N7N_A&resnum=&resdef=&showalignseq=1. (B) Align with a subset of residues 10-50 in the master chain A of PDB 1HHO. The sharable link is https://www.ncbi.nlm.nih.gov/Structure/icn3d/full.html?chainalign=1HHO_A,2HCO_A&resnum=10-50&resdef=&showalignseq=1. (C) Align with pre-defined residues: 50-100 in chain A of PDB 4N7N and residues 50-100 in chain A of PDB 1HHO. The sharable link is https://www.ncbi.nlm.nih.gov/Structure/icn3d/full.html?chainalign=1HHO_A,4N7N_A&resnum=&resdef=50-100+|+50-100&showalignseq=1.

Users can also load multiple PDB structures (either in a concatenated file or by appending PDB files with the menu “File > Open File > PDB File (appendable)”), and then use the “File > Realign Selection” feature to align multiple structures. The option “on Sequence Alignment” is similar to Figure 6B and uses a sequence alignment followed by a structure superposition. The other option “Residue by Residue” is similar to Figure 6C and uses predefined residues for a structure superposition.

### 3.6 2D Cartoons at Chain, Domain, and Secondary Structure levels

Previously, iCn3D only showed “2D Interaction” for structure data from NCBI (with the URL parameter “mmdbid=”). We recently added “2D Cartoons” for any data source such as RCSB (with the URL parameter “mmtfid=” or “pdbid=”) or EMBL-EBI (with the URL parameter “afid=”). Figure 7A shows the AlphaFold predicted structure for the UniProt ID A0A061AD48. iCn3D uses the URL parameter “afid=A0A061AD48” to get the AlphaFold predicted structure data from EMBL-EBI [1]. The high confidence parts are colored in blue and the low confidence parts are colored in yellow and orange. Figure 7B shows the 2D cartoon of the two domains in this protein: Nek and ATS1. The principal axes and size of the domains are used to determine the orientation and size of the ovals, respectively. The nodes can be dragged to modify the view of the cartoon and clicked to select in 3D.

**Figure 7.**
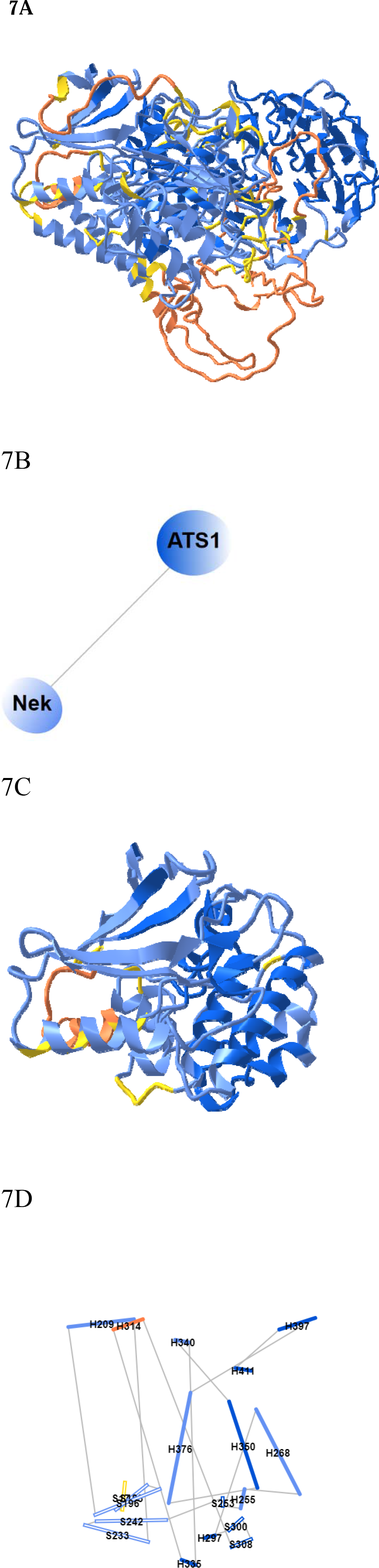
2D Cartoons of an AlphaFold structure. (A) AlphaFold predicted structure for the UniProt ID A0A061AD48. The high confidence parts of the structure are colored in blue and the low confidence parts are colored in yellow and orange. (B) 2D cartoon in the domain level. There are two domains in the protein: Nek and ATS1. (C) The Nek domain of the structure. (D) 2D cartoon at the secondary structure level. Helices are solid cylinders and labeled as “H” plus the first residue number. Sheets are empty cylinders and labeled as “S” plus the first residue number. The sharable link for panels A and B is: https://structure.ncbi.nlm.nih.gov/icn3d/share.html?R8Wtf9hDsdo6sH5Z9

Figure 7C shows the Nek domain. The 2D cartoon at the secondary structure level for the Nek domain (Figure 7D) can be shown by clicking the menu “Analysis > 2D Cartoon > Helix/Sheet Level”. The helices and sheets are projected from 3D to 2D and thus the 2D view is dynamic depending on the orientation of the 3D view. The helices and sheets can be dragged to rearrange the view and clicked to select in 3D. For a structure with multiple chains, the 2D cartoon at the chain level is useful too. The sharable link is shown in Figure 7.

### 3.7 Interactive Contact Map

iCn3D can also display interactive 2D contact maps for any subsets. Figure 8 shows the contact map of AlphaFold structure with UniProt ID A0A061AD48. The contacts within 8 angstroms of C-beta atoms are shown. The map has a scale of 0.04. If the scale is set as 1, the X- and Y-axes show residues as nodes with residue names and colors. The residue nodes and the contact points could be clicked to show the residues or contacts in 3D. This feature is available in the menu “Analysis > Contact Map”. It can be dynamically applied to multiple structures or a subset of the structure. The sharable link is shown in Figure 8.

**Figure 8.**
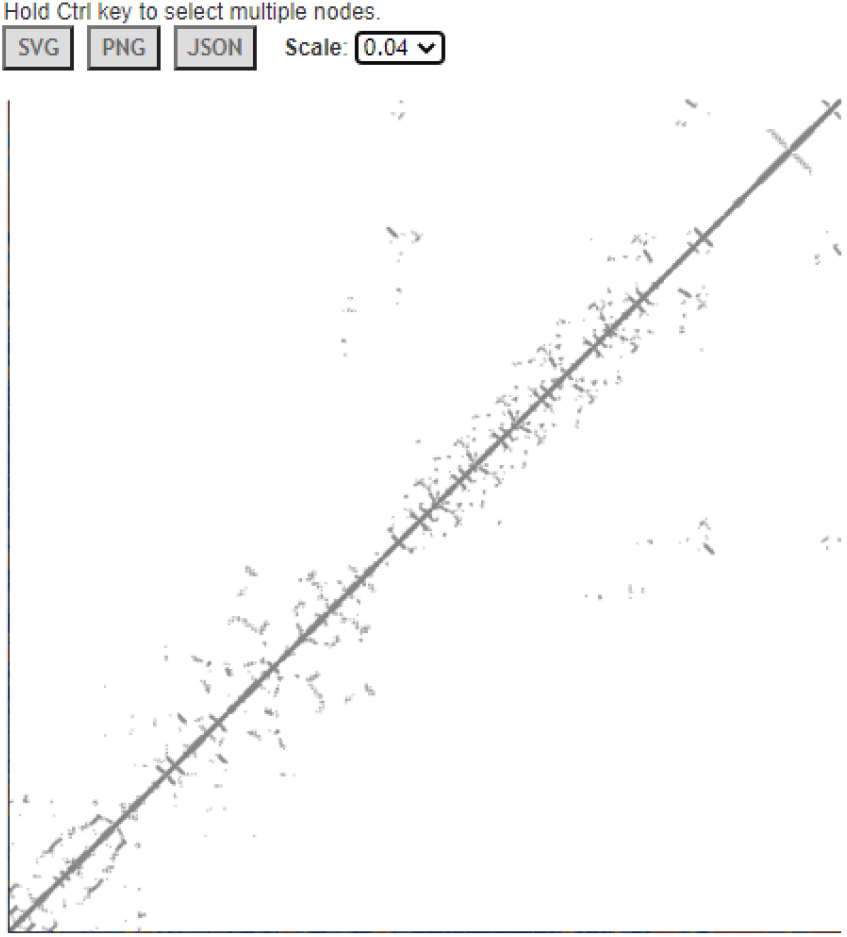
Contact map of AlphaFold structure with UniProt ID A0A061AD48. The distance threshold is 8 angstroms between C-beta atoms. The sharable link is https://structure.ncbi.nlm.nih.gov/icn3d/share.html?vTc6gkbCenTyLzt4A.

### 3.8 Python Script Based on iCn3D

Users can export files such as “iCn3D PNG Image” from iCn3D interactively. This process can be saved with a URL via the menu “File > Share Link”. Based on this URL, a simple Python script can be written to export files from iCn3D in batch mode. Some example Python scripts are listed at https://github.com/ncbi/icn3d/tree/master/icn3dpython. Users can simply replace the URL in the Python scripts to export files such as PNG images or secondary structures from iCn3D. In order to make the Python script work, users need to install “selenium”, “chrome”, “chromedriver”, and “Path” as described in the example scripts.

### 3.9 Node.js Script Based on iCn3D

While Python scripts are easy to construct as described above, they can not get all data from iCn3D. Node.js scripts can fulfil this requirement even though they are more complicated.

After we modernized iCn3D with JavaScript classes and released the npm package icn3d, we can use Node.js scripts to analyze a large set of structures in batch mode. A few Node.js scripts have been shown above in 3.1, 3.2, and 3.3. More examples are listed at https://github.com/ncbi/icn3d/tree/master/icn3dnode. Here we use the Node.js script https://github.com/ncbi/icn3d/tree/master/icn3dnode/ligand.js as an example to demonstrate the flow of the code. Users can use the command, “node ligand.js [PDB ID] [three-letter ligand residue name]”, to calculate the interactions between a ligand and a protein by calling iCn3D function “ic.showInterCls.pickCustomSphere_base”. The script first installs the required npm libraries “icn3d”, “three”, “jsdom”, and “jquery”. Then it sets up two variables used in all iCn3D functions: “me” is an instance of the iCn3DUI class, and “ic” is an instance of the iCn3D class. It then retrieves the coordinate data from NCBI and calculates the interactions. The class structure is described at https://www.ncbi.nlm.nih.gov/Structure/icn3d/icn3d.html#classstructure.

### 3.10 iCn3D in Jupyter Notebook

We released icn3dpy at https://pypi.org/project/icn3dpy/ to make iCn3D work in Jupyter Notebook. Users just need to install icn3dpy with the command “<u>pip install icn3dpy</u>” and install one lab extension with the commands “<u>pip install jupyterlab</u>” and “<u>jupyter labextension install jupyterlab_3dmol</u>”. Three commands are used to get icn3d viewer inside Jupyter Notebook: “<u>import icn3dpy</u>” to import the icn3dpy library, “<u>view=icn3dpy.view(q=‘mmdbid=6m0j’)</u>” and “<u>view</u>” to display the iCn3d viewer. The function inc3dpy.view can be optionally supplied with a command string to load a structure with a predefined view. For example, the command “<u>view = icn3dpy.view(q=‘mmdbid=6m0j’,command=‘line graph interaction pairs | !A !E | hbonds, salt bridge, interactions, halogen, pi-cation, pi-stacking | false | threshold 3.8 6 4 3.8 6 5.5; show selection; add residue number labels; set label scale 2.0|||{“factor”:“0.6758”,“mouseChange”:{“x”:“0.000”,“y”:“0.000”},”quaternion”:{“_x”:“-0.6336”,“_y”:“0.3127”,“_z”:“-0.3564”,“_w”:“0.6114”}}’)</u>” produces the view as in Figure 2. The string “q=‘mmdbid=6m0j’” specifies the structure to be loaded and the command string after the string “command=” is used to set the predefined display. This command string can be retrieved from a web session. It is the string after the string “command=” in the expanded URL of https://structure.ncbi.nlm.nih.gov/icn3d/share.html?oLwZzGL59izJeEBVA from Figure 2.

## 4 Discussion

Users can use either Python scripts or Node.js scripts to a large set of 3D structures. While the Python scripts are easy to construct, the iCn3D-based Node.js scripts fully leverage the power of iCn3D and can achieve any feature in iCn3D. We’ll add more export features in iCn3D so that users can easily generate Python scripts to export files from iCn3D.

There are many ways to save work in iCn3D such as shortened URLs, iCn3D PNG images, or state files, etc. The recommended way is to save an iCn3D PNG image with the menu “File > Save File > iCn3D PNG Image > Original Size & HTML”. Both an image and an HTML file are saved. The iCn3D PNG image not only contains the custom view, but also contains all data and the command history. It can be loaded back to iCn3D with the menu “File > Open File > iCn3D PNG Image”. It can also be stored in a web server, and be retrieved to reproduce the saved views, e.g., https://www.ncbi.nlm.nih.gov/Structure/icn3d/full.html?type=icn3dpng&url=https://www.ncbi.nlm.nih.gov/Structure/icn3d/pdb/3GVU.png. The saved HTML files can be concatenated to form a gallery page, similar to the iCn3D gallery page at https://www.ncbi.nlm.nih.gov/Structure/icn3d/icn3d.html#gallery.

We showed three examples to convert software packages to RESTful APIs so that iCn3D can use them in AJAX calls seamlessly. To follow the FAIR (Findable, Accessible, Interoperable and Reusable) guiding principles [13; 14], bioinformatic tools had better provide RESTful APIs to make these tools accessible and reusable.

Our future plan includes linking iCn3D with multiple sequence alignment tools, adapting WebGL2 or WebGPU in iCn3D, and more.

## Conflict of Interest

*DNAnexus Inc. and Noma Technology were affiliated with this study. The authors declare that the research was conducted in the absence of any commercial or financial relationships that could be construed as a potential conflict of interest*.

## Author Contributions

Philippe Youkharibache is the driving force for the development of iCn3D. He provides many designing ideas and consistently tests the features of iCn3D. As part of iCn3D team, Aron Marchler-Bauer, Christopher Lanczycki and Gabriele Marchler give many suggestions and test iCn3D features. Dachuan Zhang, Shennan Lu and Thomas Madej provide data for iCn3D from backend servers. Tiejun Cheng developed the Python code to export 3D images in batch mode. The following people contributed to the iCn3D feature of 2D cartoon during the ISMB Codeathon 2021: Li Chuin Chong, Sarah Zhao, Kevin Yang, Jack Lin, and Zhiyu Cheng. The following people contributed to the iCn3D features of dynamic symmetry and side chain prediction due to mutation during the STRIDES Codeathon 2020: Rachel Dunn, Sridhar Acharya Malkaram, Chin-Hsien Tai, and David Enoma. The following people contributed to using iCn3D in Jupyter Notebook during the ISMB Codeathon 2020: Ben Busby, Nicholas Johnson, Francesco Tabaro, Guangfeng Song, and Yuchen Ge.

## Funding

The work of JW, PY, AM, CL, DZ, SL, TM, GHM, TC, and GS was supported by the National Center for Biotechnology Information of the National Library of Medicine (NLM), National Institutes of Health. Comments, suggestions and questions are welcome and should be directed to. Funding to pay the Open Access publication charges for this article was provided by the National Center for Biotechnology Information of the National Library of Medicine (NLM), National Institutes of Health.

## Acknowledgments

We would like to specially thank Barry Honig for licensing DelPhi and providing scap to us; Emil Alexov for his very helpful suggestions about using DelPhi; David Koes for his help in using iCn3D in Jupyter Notebook; Ravinder Abrol, Raul Cachau, Todd Smith and Sandra Porter for their feedback and suggestions during the development of iCn3D; and Ravinder Abrol and Allissa Dillman for organizing several codeathons related to iCn3D.

## Notes

### Competing Interest Statement

The authors have declared no competing interest.

